# Designing Experimental Protocols to Elicit Vigilance Variations in Controlled Laboratory Settings

**DOI:** 10.1101/2024.10.01.616166

**Authors:** Saeed Zahran

## Abstract

Vigilance plays a vital role in numerous high-risk professions, where real-time monitoring of vigilance levels is highly beneficial. Brain-Computer Interfaces (BCIs) utilizing EEG signals, along with machine learning and deep learning algorithms, present a promising solution for the classification and monitoring of vigilance levels. This study aimed to create a dataset for training such models by evaluating four experimental paradigms: the Hitchcock Air Traffic Controller Task (ATC), the simultaneous line task, the successive line task, and the Oddball task. Subjective reports and behavioral performance were analyzed to determine the effectiveness of these tasks in inducing vigilance decrements over time. The findings reveal that both the ATC and simultaneous line tasks effectively induced significant declines in subjective vigilance ratings and behavioral performance. In contrast, the Oddball task was less successful in generating a noticeable vigilance decrement. This research demonstrates the potential of the ATC and simultaneous line tasks to induce vigilance variations, providing valuable datasets for training vigilance detection algorithms. Additionally, it highlights the importance of considering non-linear fluctuations in vigilance and the need for more advanced techniques to accurately classify different vigilance states. Such improvements in vigilance monitoring could substantially enhance safety and well-being in critical work environments.

## I. Introduction

This Vigilance refers to the sustained ability to maintain attention and focus on a task over an extended period [1, 2]. It plays a critical role in everyday life, influencing task performance, safety, and decision-making. In numerous professions, maintaining high levels of vigilance is essential but can also be quite challenging. For example, technicians at ENEDIS, the public electricity distribution company in France, must remain highly vigilant throughout their workday, often operating in dangerous environments such as high-voltage electrical stations. Any lapse in attention in these contexts could have life-threatening consequences. Other professions that require continuous vigilance include air traffic controllers, long-distance drivers, security personnel, and baggage screeners [3]. Engaging in prolonged tasks often leads to a decline in vigilance over time, a phenomenon known as “vigilance decrement” [4, 5, 6]. In laboratory settings, this decrement typically manifests as a reduction in the correct detection of infrequent targets, an increase in false alarms, and slower reaction times [7, 8, 9]. This decline usually becomes apparent within the first 20-35 minutes of task engagement [10, 11]. Vigilance decrements are particularly noticeable in monotonous tasks with few critical events, both in laboratory environments and in real life. Even though these tasks may seem repetitive, missing a critical target can have severe consequences. The ability to assess and monitor an individual’s vigilance state in real-time holds significant potential for improving safety and overall worker well-being. One promising approach for real-time vigilance monitoring is the use of Brain-Computer Interfaces (BCIs). BCIs measure brain activity, typically through electroencephalography (EEG), to extract features or biomarkers, which are then classified using machine learning or deep learning algorithms. In the context of vigilance detection, BCIs could provide warnings for critical drops in vigilance. Recent years have seen increasing efforts to classify vigilance levels based on EEG signals obtained in laboratory settings [3, 11,12,13].

Training models to accurately detect varying levels of vigilance using EEG data requires large datasets of physiological signals, accompanied by corresponding labels (e.g., “high” vs. “low” vigilance or scales of vigilance). As obtaining and accurately labeling EEG data in real-world work environments can be challenging, most datasets are collected in controlled laboratory settings.

The primary aim of this study was to identify effective laboratory paradigms that could induce decrements or fluctuations in vigilance. By gathering physiological data from participants performing these tasks, we could create a dataset suitable for training models to classify different vigilance states. Ultimately, this work aims to develop an online vigilance monitoring system for real-world applications, such as for ENEDIS workers.

Through a review of the literature, we identified four experimental paradigms that appeared particularly promising for reliably inducing vigilance decrements. The following sections will provide a detailed description of these paradigms, followed by an analysis of the behavioral outcomes and subjective reports from participants. The appropriateness of these experimental tasks will be discussed in light of the results.

## II. MATERIALS AND METHODS

### Vigilance Tasks

#### Hitchcock Air Traffic Controller Task (ATC)

The Hitchcock Air Traffic Controller Task (ATC), first proposed by Hitchcock et al. (2003) [14], has been widely adopted in subsequent vigilance studies [11, 15-17]. This task simulates air-traffic control through a radar display featuring three concentric white rings on a black background, with a red dot in the center representing a city. Two airplanes are depicted as white lines, which can be oriented either in a northeast-southwest or northwest-southeast direction. In the critical condition, the planes are positioned directly opposite each other, signaling a potential collision, requiring the participant to intervene. In the non-critical condition, the planes are slightly misaligned, indicating no immediate danger. Participants respond to critical signals by pressing the “Enter” key on a keypad.

In this study, we adapted the task using stimulus parameters consistent with those from previous research: the radii of the white circles were 1.8, 3, and 4.2 cm, with a thickness of 0.75 mm. The red dot had a radius of 5.25 cm and was surrounded by an additional white circle with a 6 cm radius and a 0.75 mm thickness. The airplane lines were 3 cm in length and nearly touched the innermost ring. In the non-critical condition, the lines were offset by 8 mm horizontally.

Each stimulus was presented for 300 milliseconds, with 2-second intervals between stimuli. The experiment included 1,050 stimuli and lasted 40 minutes without breaks. The critical events had a probability of 3.3%, resulting in approximately 35 target stimuli. These critical events were evenly distributed throughout the session by dividing the task into four segments of equal duration. Each segment contained an equal number of critical stimuli, ensuring they did not occur consecutively.

Before the experimental block, participants completed a training block of 200 stimuli to familiarize themselves with the task and minimize learning effects during the experimental session. Feedback regarding correct responses, hits, misses, and false alarms was provided every 50 trials.

#### Line Tasks (Simultaneous & Successive)

The line tasks, developed by Parasuraman and Mouloula (1987) [18], required participants to discriminate between pairs of vertical lines displayed for 150 milliseconds (Figure 2). We utilized two versions of this task: in the simultaneous discrimination task, critical stimuli consisted of pairs where one line was shorter than the standard length, with the shorter line randomly assigned to either the left or right side. In the successive discrimination task, both lines in the critical stimuli were shorter than the standard lines.

**Figure 1:**
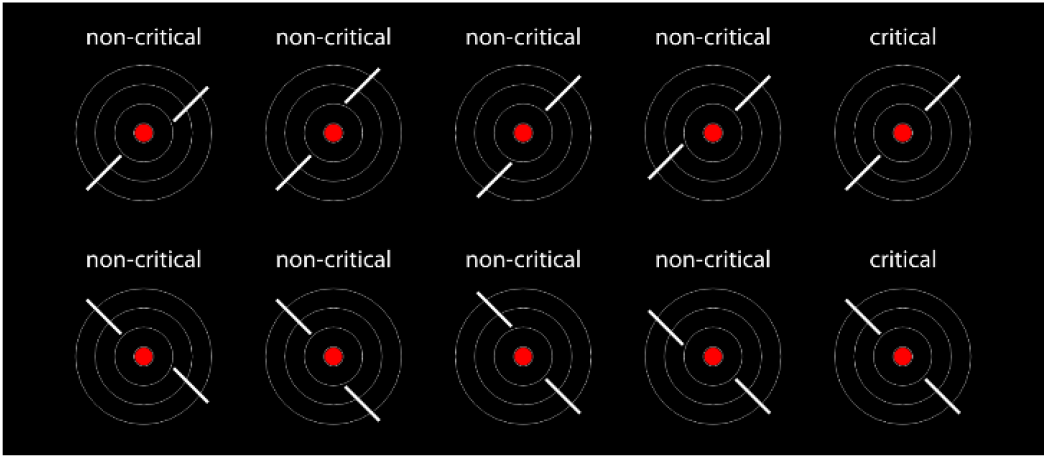
Critical and non-critical stimuli of the Air Traffic Controller Task.

**Figure 2:**
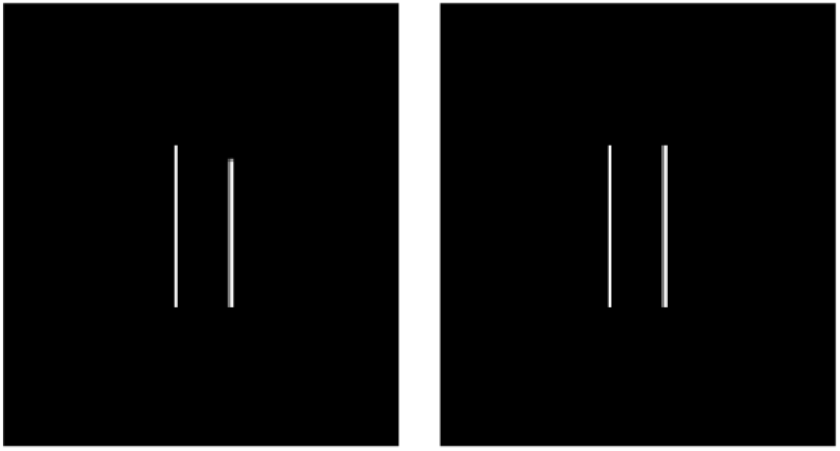
Line task. Left: Example for a target stimulus of the simultaneous version, with unequal line length. Right: Noncritical stimulus.

In the original study, line length differences were varied to adjust the task difficulty. In our version, we used a standard line length of 3.6 cm, with a difference of 1.2 mm in the simultaneous task and 3.4 mm in the successive task. The width of the lines was set at 7.5 mm. The experiment included 1,460 trials, with a critical stimulus probability of 10%. As in the ATC task, critical stimuli were evenly distributed throughout the task by dividing the block into four equal segments.

Participants first completed a brief training session of 10 trials, followed by feedback after each trial. A longer practice session of 200 trials followed, with feedback on the percentage of correct responses, as well as the number of hits, misses, and false alarms provided every 50 trials.

#### Oddball Task

The 3-stimulus Oddball task, adapted from Comerchero et al. (1999) [19], involved three different visual stimuli: targets, non-target distractors, and standard distractors (Figure 3). Targets and non-target distractors were presented in 10% of the trials. Targets had a radius of 1.25 cm, while standard distractors had a radius of 0.7 cm. Non-target distractors measured 1.4 x 1.4 cm. Participants completed 20 practice trials without feedback, followed by four blocks of 300 trials each. Feedback was provided at the end of each block.

**Figure 3:**
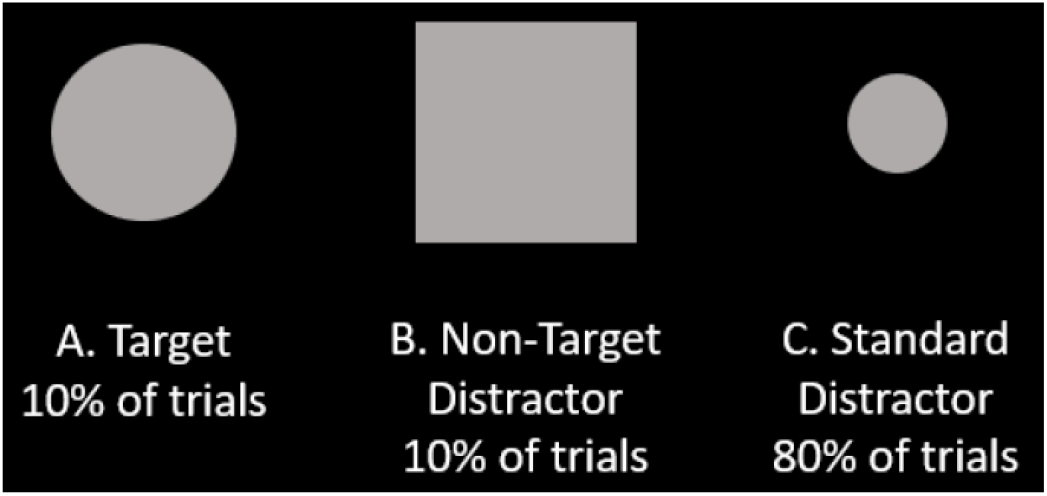
Oddball task stimuli. Taken from Kamrud et al. 2021.

### Questionnaires

Two questionnaires were used to measure vigilance and workload. Before the experiment, participants completed the Stanford Sleepiness Scale (SSS), a 7-point Likert scale assessing subjective sleepiness [20]. Participants selected from seven statements describing their current level of alertness, ranging from “Feeling active and vital, alert, wide awake” to “Almost in reverie, sleep onset soon, lost struggle to remain awake.” A sleepiness score between 0 and 6 was assigned based on the selected statement. Participants were also asked to report the number of hours they had slept the previous night.

After the experiment, participants completed the NASA-Task Load Index (NASA-TLX), a questionnaire designed to assess workload [21]. The original version includes six questions rated on a 20-point Likert scale, evaluating mental demand, physical demand, temporal demand, subjective performance, effort, and frustration. In this study, four additional questions were included: 1) “How stressful was the task?”; 2) “How difficult was it for you to stay awake and concentrated?”; “What was your subjective vigilance level during the first half of the task?”; and 4) “What was your subjective vigilance level during the second half of the task?” These questions aimed to gauge how effectively the tasks induced drops in subjective vigilance.

### Participants

Thirty-two subjects (13 female, 19 male) participated in the study. One participant completed all four tasks, two participants completed three tasks, five participants completed two tasks, and the remaining participants completed one task each. The age range was 20 to 60 years, with a mean age of 34.1 ± 10 years.

### Behavioral Performance Indices

Several behavioral performance indices were calculated. Accuracy was defined as the proportion of correct responses out of all trials:

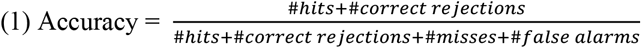

Reaction time (RT) was computed only for correct selections:

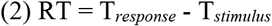

To combine both speed and accuracy into a single performance measure, two methods were employed. First, the Inverse Efficiency Score (IES) [Townsend & Ashby, 1978, 1983] was calculated:

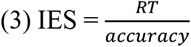

A lower IES indicates better performance. Second, the Balanced Integration Score (BIS), a z-score combining accuracy and RT, was calculated [22]. Accuracy and RT were standardized as follows:

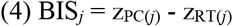

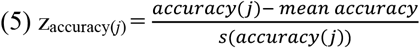

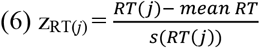

Higher BIS scores indicate higher performance.

### Statistics

Mixed-effects linear regression analyses were used to compare task types while accounting for individual differences among participants, as some completed multiple tasks. Likelihood ratio tests (LRTs) were conducted to determine whether the inclusion of task type significantly improved model fit.

Paired sample t-tests were used to compare vigilance ratings between the first and second halves of the experiment. Cohen’s d effect size was calculated using the formula:

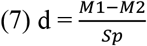

Where M1 and M2 are the means of the two groups, and S_p is the pooled standard deviation. Effect sizes were interpreted as follows:

- 0.2 = small effect
- 0.5 = moderate effect
- 0.8 = large effect

## III. RESULTS

### Questionnaires

On average, participants slept for 6.9±1 hours the night before the experiment (range: 5-9 hours), with a mean sleepiness score of 1.3±0.9 (range: 0-4). Mental workload was rated at 12.6±6.4 on average (range: 1-20). A significant effect of task on mental workload was indicated by the LRT (L.-ratio = 12.19, p = .0067), with the highest mental demand observed in the successive line task (15.83±2.85), followed by the simultaneous line task (13±6.54), the ATC task (12.59±5.77), and the Oddball task (6.8±6.4) (Figure 4, top left). Physical workload was rated at 7.6±6 (range: 1-20), with no significant effect of task (L.-ratio = 2.45, p = .48).

**Figure 4.**
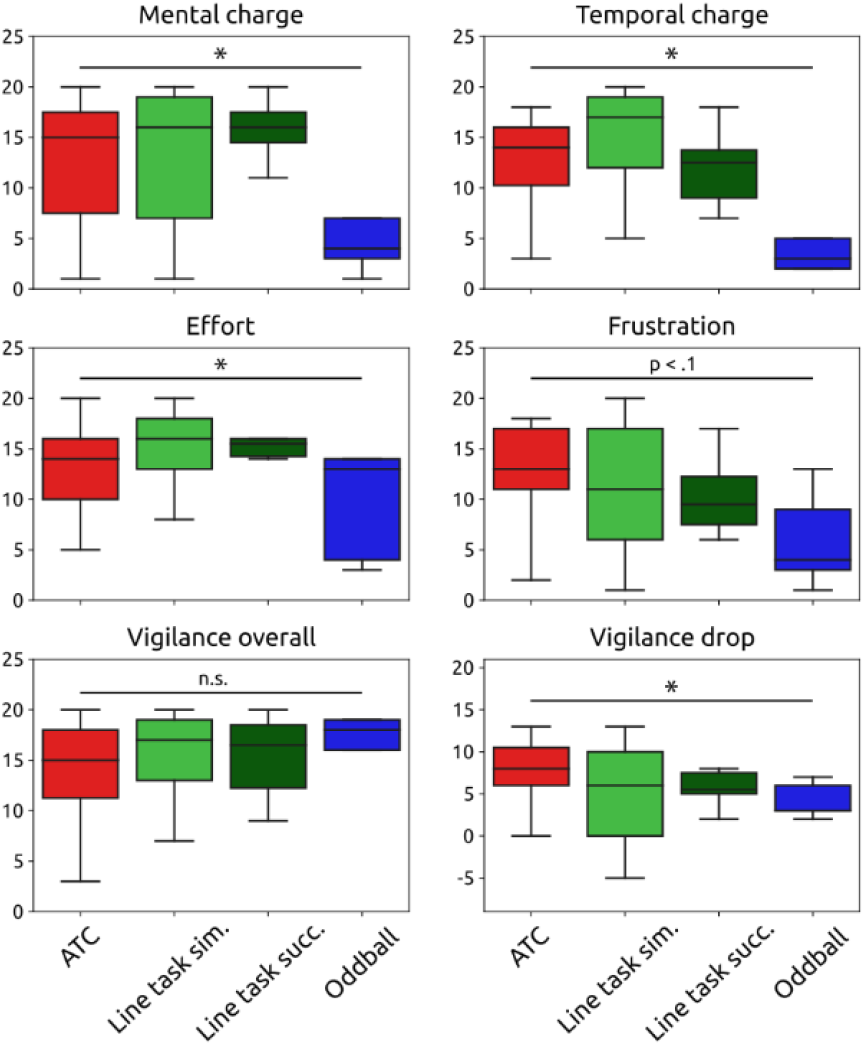
Scores from the NASA questionnaire for selected questions

**Figure 5.**
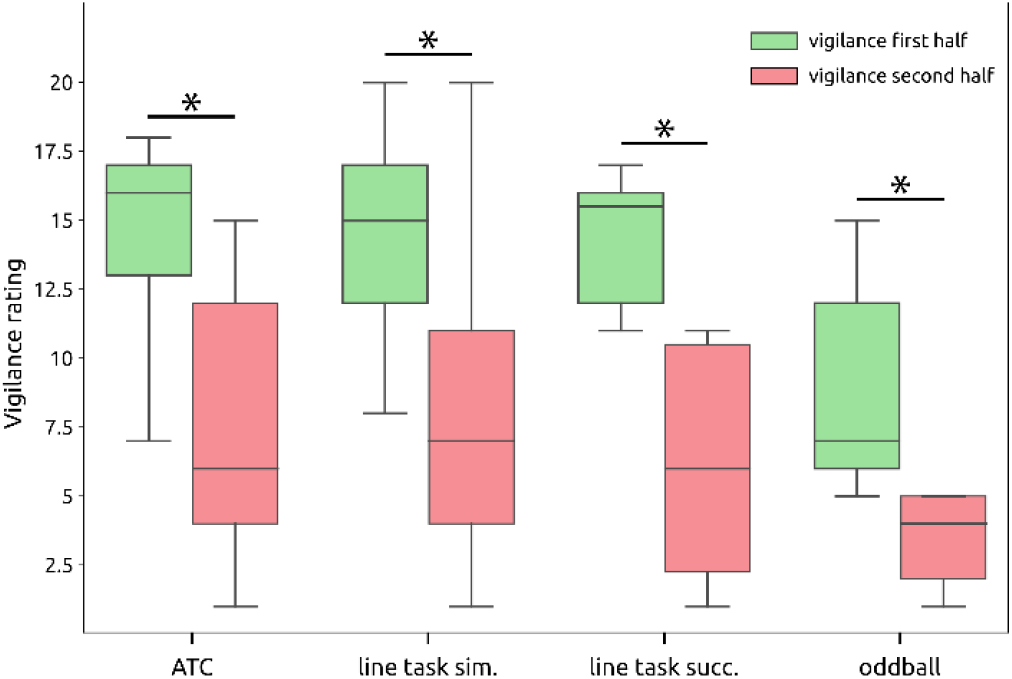
Vigilance ratings for the first vs. second half of the block, by task. All tasks showed significant drops in subjective vigilance.

Temporal demand averaged 10.3±4.8 (range: 1-18), and LRT analyses revealed a significant effect of task (L.-ratio = 24.26, p < .0001). The highest temporal demand was reported for the simultaneous line task (15.29±4.11), followed by the ATC task (13.02±4.13), the successive line task (12±3.7), and the Oddball task (4.8±3.76) (Figure 4, top right).

Subjective performance ratings averaged 10.3±4.8 (range: 1-18), with no significant effect of task (L.-ratio = 2.25, p = .52).

Effort was rated at 13.5±4.6 (range: 3-20) and showed a significant effect of task (L.-ratio = 9.1, p = .03), with the highest effort reported for the successive line task (15.17±2.97), followed by the simultaneous line task (14.76±4.26), the ATC task (12.93±4.36), and the Oddball task (9.6±5) (Figure 4, middle left).

Frustration averaged 11.5±5.8 (range: 1-20), showing a trend toward a task effect (L.-ratio = 7.34, p = .06). The highest frustration was reported for the ATC task (13±4.82), followed by the simultaneous line task (11.38±6.49), the successive line task (10.33±3.73), and the Oddball task (6±4.38) (Figure 4, middle right).

Stress was rated at 6.7±4.6 (range: 1-17) with no significant effect of task (L.-ratio = 3.18, p = .37).

Overall vigilance during the tasks was rated at 14.8±4.9 (range: 2-20), with no significant effect of task (L.-ratio = 0.13, p = .99) (Figure 4, bottom left).

To assess the drop in vigilance, we calculated the difference in vigilance ratings between the first and second halves of the experiment, with positive values indicating a greater reduction in vigilance during the second half. The mean vigilance drop was 6.7±5.2 (range: -8-16) and showed a significant task effect (L.-ratio = 10.56, p = .01). The ATC task produced the largest vigilance decrement (7.54±4.72), followed by the successive line task (7±4.4), the simultaneous line task (5.53±5.34), and the Oddball task (4.2±1.94) (Figure 4, bottom right).

Paired t-tests showed significantly higher vigilance ratings during the first half of the task compared to the second half for the ATC task (t(50) = 7.5, p < .0001, Cohen’s d = 1.43), the simultaneous line task (t(50) = 4.14, p < .001, Cohen’s d = 1.05), the successive line task (t(50) = 3.59, p = .02, Cohen’s d = 1.33), and the Oddball task (t(4.33) = p = .01, Cohen’s d = 0.8). The largest effect size was observed for the ATC task, followed by the successive line task, the simultaneous line task, and the Oddball task.

In summary, mental and temporal demand, as well as effort, varied significantly across tasks, with the Oddball task consistently requiring the least. Although overall vigilance did not differ significantly between tasks, the decline in subjective vigilance was most pronounced for the ATC task and the two line tasks and least for the Oddball task.

### Performance

#### Overall and Comparison Between Tasks

The mean hit rate was 0.96±0.24, with a significant task effect (L.-ratio = 24.26, p < .0001). The highest hit rate was observed for the Oddball task (0.91±0.14), followed by the ATC task (0.82±0.13), the simultaneous line task (0.55±0.22), and the successive line task (0.42±0.22) (Figure 6, top left).

**Figure 6.**
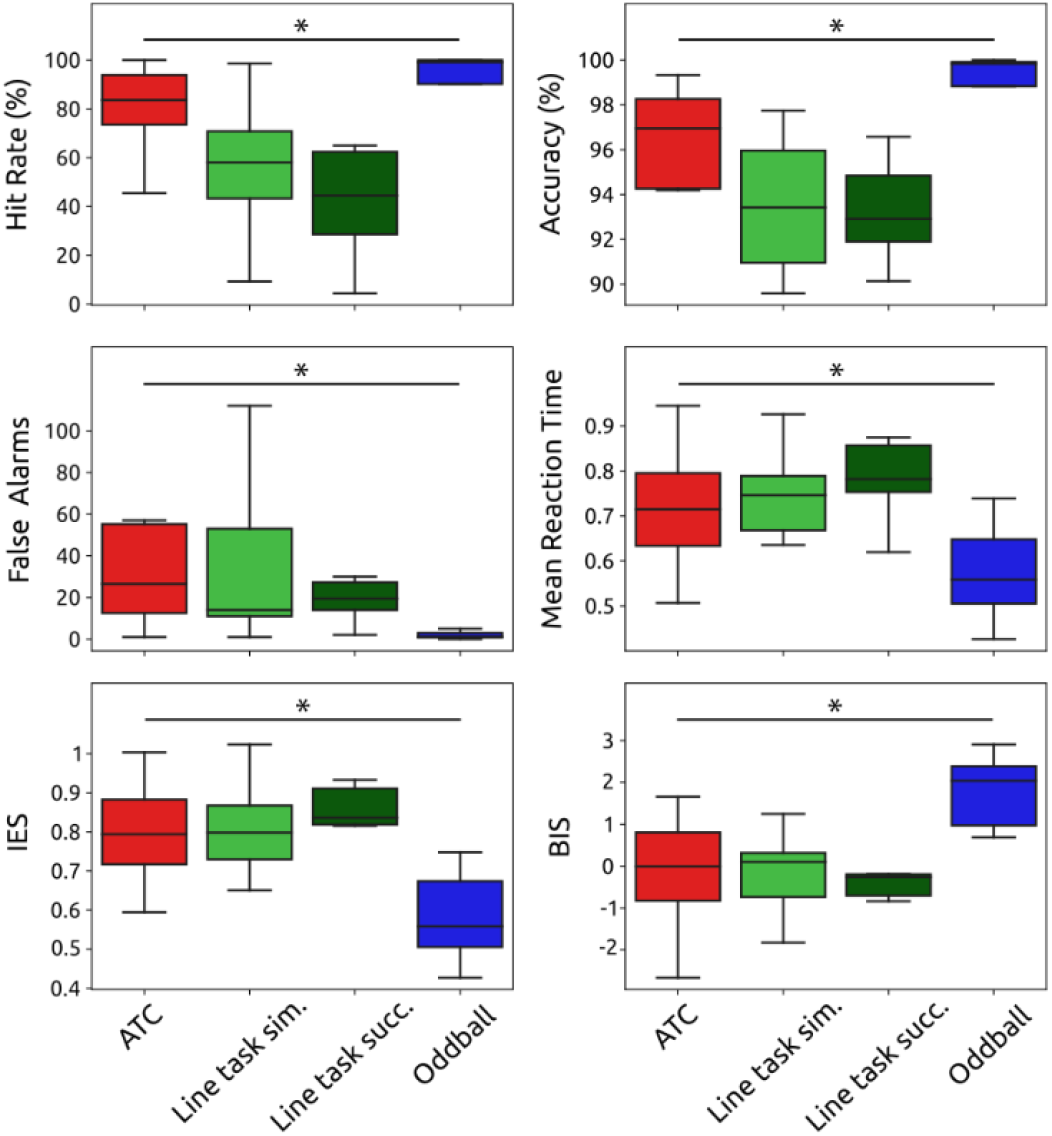
Behavioral performance measures for each task. BIS: Balanced Integration Score; IES: Inverse Efficiency Score.

Mean accuracy was 0.93±0.08, with no significant task effect (L.-ratio = 3.54, p = .32). The mean number of false alarms was 55.9±103.09, also showing no significant task effect (L.-ratio = 3.27, p = .35).

Mean reaction time (RT) was 0.73±0.15 seconds, with significant variation between tasks (L.-ratio = 17.9, p < .001). The fastest RT was recorded for the Oddball task (0.58±0.11 seconds), followed by the ATC task (0.73±0.12 seconds), the simultaneous line task (0.75±4.11 seconds), and the successive line task (0.83±0.17 seconds) (Figure 6, middle right).

Mean Inverse Efficiency Score (IES) was 0.79±0.16, with a significant task effect (L.-ratio = 20.46, p < .001). The Oddball task showed the lowest IES (0.58±0.12), followed by the ATC task (0.79±0.12), the simultaneous line task (0.82±0.15), and the successive line task (0.89±0.2) (Figure 6, lower left).

The mean Balanced Integration Score (BIS) was 0±1.27 and showed a significant task effect (L.-ratio = 16.67, p < .001). The highest BIS was recorded for the Oddball task (1.8±0.84), followed by the ATC task (0±1.1), the simultaneous line task (-0.32±1.09), and the successive line task (-0.59±1.32) (Figure 6, lower right).

#### Vigilance Decrement

To examine whether the tasks induced performance decrements over time, we divided the 40-minute task into four bins of 10 minutes each and calculated performance for each bin. Figure 7 illustrates the average performance for each measure, task, and time bin. Hit rate appeared to decline slightly over time for the ATC task, simultaneous line task, and successive line task. Accuracy decreased for the ATC task and the successive line task. RT slowed over time for the ATC task and, to a lesser extent, the simultaneous line task. The ATC and simultaneous line tasks showed a marked increase in false alarms. IES increased significantly for the ATC task and slightly for the simultaneous line task, while it remained stable for the Oddball task and decreased for the successive line task. BIS showed a decline over time for the ATC task, but no significant changes were observed in the other tasks.

**Figure 7.**
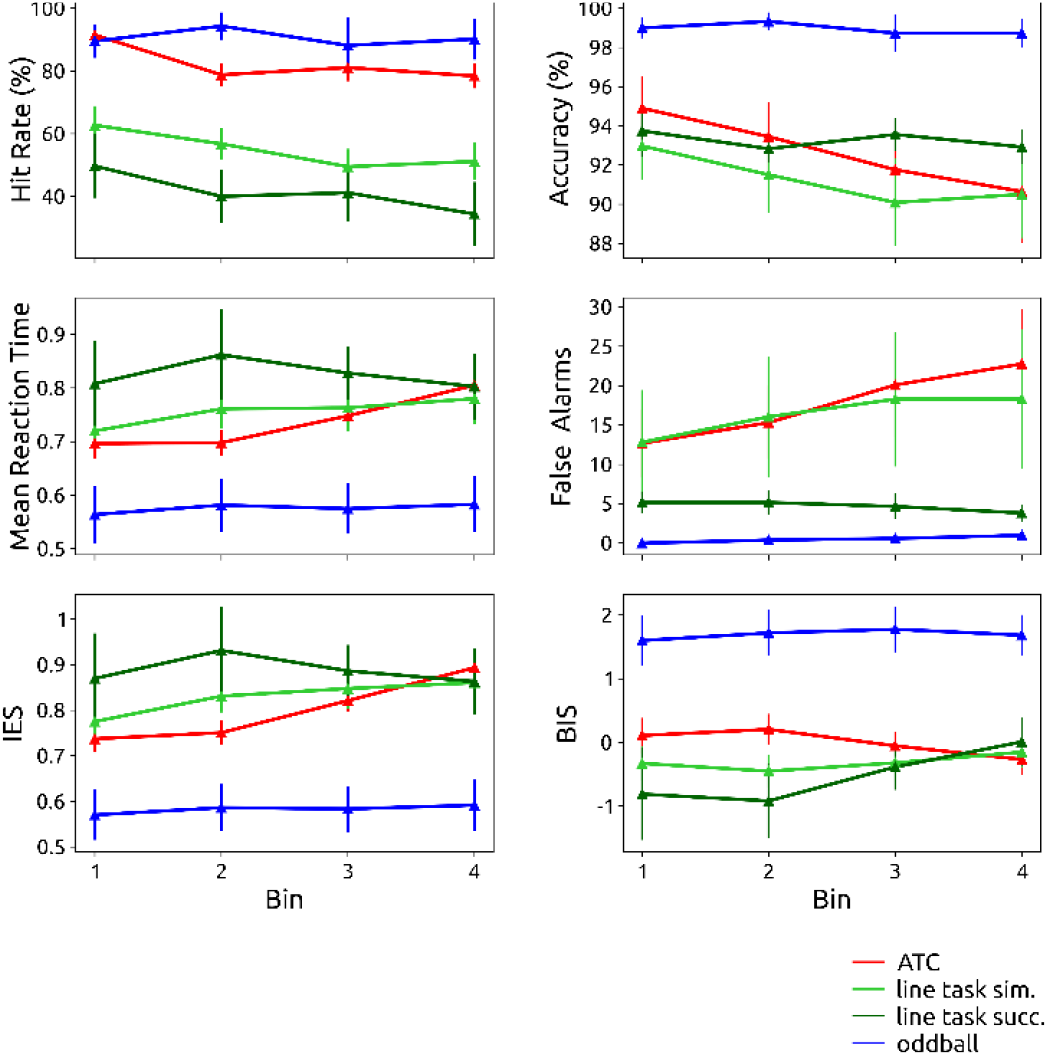
Evolution of behavioral performance over bins for each performance measure and bin. BIS: Balanced Integration Score; IES: Inverse Efficiency Score.

Overall, the vigilance decrement was not consistently observed across all measures and tasks. The ATC and simultaneous line tasks demonstrated the clearest declines in performance across several metrics.

Figure 8 illustrates individual patterns of performance evolution, as represented by IES. While the mean IES tended to increase over time, individual participants showed varying patterns: some participants exhibited a steady decline, others showed initial declines followed by recovery, and some alternated between declining and improving performance. However, the ATC and simultaneous line tasks were associated with the most consistent increases in IES among participants.

**Figure 8.**
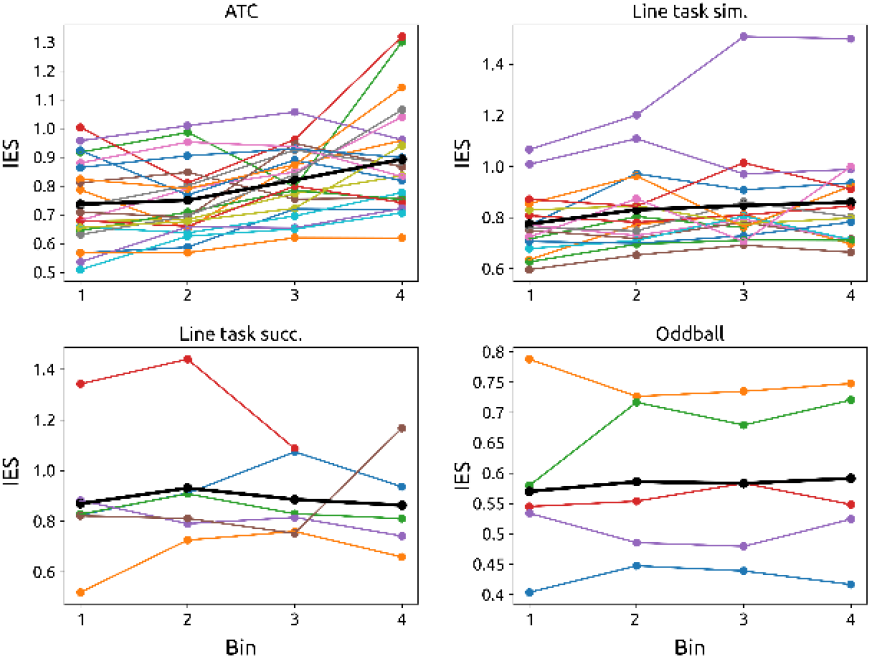
Inverse Efficiency Score by task, bin and subject. Each colored line represents one subject. Black lines: Mean IES for task.

## Discussion

This study evaluated the effectiveness of four vigilance tasks—the ATC task, simultaneous line task, successive line task, and Oddball task—in inducing vigilance decrements, as reflected in both subjective ratings and behavioral performance. From the questionnaire data, we observed notable differences in the mental and temporal demands, as well as the effort required for each task, with the Oddball task consistently presenting the lowest demand. While all tasks led to a decline in vigilance from the first to the second half of the experiment, the ATC task and the two line tasks exhibited the most pronounced drops in perceived vigilance, whereas the Oddball task showed the least decline.

In terms of behavioral performance, participants performed best on the Oddball task, followed by the ATC task and the line tasks. The most substantial behavioral vigilance decrements were observed during the ATC and simultaneous line tasks, while the successive line task and the Oddball task exhibited less of a decrement.

These findings suggest that the ATC and simultaneous line tasks are the most suitable for inducing vigilance drops in a controlled setting. The Oddball task, by contrast, appears less effective, likely due to its lower task demands. Previous research has demonstrated that vigilance decrement is more pronounced and occurs more quickly in tasks with higher cognitive demands [23]. Thus, the Oddball task may not have been challenging enough to significantly deplete participants’ cognitive resources over time.

However, even in the ATC task, the behavioral performance did not show a linear vigilance decrement for all participants. Some individuals exhibited recovery between time bins, or alternating periods of low and high performance. This non-linear pattern suggests that vigilance may fluctuate randomly rather than decline steadily. For instance, a participant struggling to stay awake might occasionally recover temporarily. This variability complicates the process of accurately labeling segments of the task with distinct vigilance levels. Within the chosen 10-minute bins, participants may have experienced multiple phases of higher and lower vigilance. The bin size is arbitrary, and vigilance fluctuations may occur across various time scales, making it difficult to establish a “ground truth” for vigilance using behavioral performance alone. Biomarkers such as cortisol levels or pupil diameter might provide more reliable indicators of vigilance levels. Alternatively, more advanced computational methods, such as unsupervised learning techniques, could be explored to detect vigilance states more accurately [24].

A practical solution to address this issue is to focus on data from participants who exhibited clear-cut vigilance decrements, as indicated by both the questionnaires and behavioral performance. Moreover, despite variability in performance over time, most participants showed reduced performance toward the end of the experiment compared to the beginning. As such, data from the beginning of the experiment can be used to represent “high vigilance” states, while data from the end can represent “low vigilance” states. This approach was similarly employed by Kamrud et al. [11], though it does result in some loss of data.

Ultimately, it is unlikely that any task will reliably induce a purely linear decrease in vigilance across all participants. However, this study has demonstrated that the ATC task and the line tasks are effective in inducing decreases in vigilance over time, making them suitable for generating different vigilance states in a laboratory setting. These tasks could play a key role in the development of vigilance monitoring systems, which have the potential to significantly enhance safety and well-being in critical work environments.

